# Small cell lung cancer co-culture organoids provide insights into cancer cell survival after chemotherapy

**DOI:** 10.1101/2023.01.03.522668

**Authors:** Chandani Sen, Caroline Koloff, Souvik Kundu, Dan C Wilkinson, Juliette Yang, David W Shia, Luisa K Meneses, Tammy M Rickabaugh, Brigitte N Gomperts

**Affiliations:** UCLA Children’s Discovery and Innovation Institute, Mattel Children’s Hospital, Department of Pediatrics, David Geffen School of Medicine, UCLA, Los Angeles, CA, USA, 90095; Pulmonary Medicine, David Geffen School of Medicine, UCLA, Los Angeles, CA, USA, 90095; Jonsson Comprehensive Cancer Center, UCLA, Los Angeles, CA, USA, 90095; Eli and Edythe Broad Stem Cell Research Center, UCLA, Los Angeles, CA, USA, 90095; Intel Labs, San Diego, CA, USA, 92127

**Keywords:** small cell lung cancer, three-dimensional organoid model, cancer-stromal interaction, cancer relapse, automated image analysis

## Abstract

Small-cell-lung-cancer (SCLC) has the worst prognosis of all lung cancers because of a high incidence of relapse after therapy. We developed a bioengineered 3-dimensional (3D) SCLC co-culture organoid as a phenotypic tool to study SCLC tumor kinetics and SCLC-fibroblast interactions during relapse. We used functionalized alginate microbeads as a scaffold to mimic lung alveolar architecture and co-cultured SCLC cell lines with primary adult lung fibroblasts (ALF). We found that SCLCs in the model proliferated extensively, invaded the microbead scaffold and formed tumors within just 7 days. We compared the bioengineered tumors with patient tumors and found them to recapitulate the pathology and immunophenotyping of the patient tumors better than the PDX model developed from the same SCLC cell line. When treated with standard chemotherapy drugs, etoposide and cisplatin, the organoid recapitulated relapse after chemotherapy. Co-culture of the SCLC cells with ALFs revealed that the fibroblasts play a key role in inducing faster and more robust SCLC cell regrowth in the model. This was a paracrine effect as conditioned medium from the same fibroblasts was responsible for this accelerated cell regrowth. This model is also amenable to high throughput phenotypic or targeted drug screening to find new therapeutics for SCLC.

## Introduction

Lung cancer is the largest cause of cancer death in both men and women in the United States, with an overall 5-year survival rate of only about 25%, according to the Surveillance, Epidemiology, and End Results (SEER) data [1]. However, this overall 5-year survival rate decreases to 8% if distant tumor spread is present at diagnosis. Of all the lung cancer subtypes, small cell lung cancer (SCLC), a neuroendocrine sub-type representing about 10% of all lung cancers, has by far the worst prognosis and is often highly metastatic [2–4]. The classic neuroendocrine markers (synaptophysin, chromogranin A, and NCAM) are usually used for diagnosis [5] of the tumor biopsy sample and the cisplatin-etoposide combination is used as standard first line chemotherapy [4]. The main cause of death in these patients is resistance to chemotherapy as most patients respond well to initial therapy but will experience tumor recurrence within one year after completing chemotherapy [6]. The underlying pathogenesis of this tumor resistance is still unknown and there have been no advances in therapies for more than three decades [7].

Drug screening for cancer is usually performed initially with cancer cell lines grown in aggregates in plastic dishes. These cells are easy and cheap to grow and convenient for high throughput screening but they fail to represent the complexity of the tumor microenvironment [8]. These kinds of cancer models show about a 10% success rate in developing anti-cancer drugs for clinical trials. But because it is challenging to model relapse in SCLC, most of the drugs identified failed at preventing recurrence in pre-clinical and clinical trials [9]. The current gold standard patient-derived xenografts (PDX), where cancer cell lines or small pieces of tumor tissue derived from patients are transferred to immune-deficient mice, also has several limitations, including chances of tumor tissue engraftment failure, a long tumor development timeline, dissimilarity of the tumor microenvironment between human and murine models, and low throughput for drug screening [10]. Replacement of animal PDX modeling will also address the 3Rs’ (replacement, refinement and reduction) that are essential for animal welfare [26]. Recently, 3D human organoid/spheroid models are gaining popularity in cancer research [11–14], but currently there is no existing human co-culture lung organoid model to study the tumor microenvironment and phenotypic changes after chemotherapy that can be useful in drug screening for cancer relapse.

To address these issues, we have developed a scaffold-based 3D organoid mimicking lung micro-architecture using primary human healthy adult lung fibroblasts (ALF) and SCLC cell lines. This co-culture organoid shows phenotypic changes consistent with disease progression *in vitro* and allows the assessment of the role of the fibroblasts in the development of disease recurrence after chemotherapy. In addition, our model is scalable for 96-well and 384-well plates and therefore is valuable for high throughput screening (HTS) for therapeutic strategies to prevent SCLC relapse.

## Results

### The co-culture SCLC organoid recapitulates tumor growth, local invasion and the mechanobiology of SCLC

To recapitulate the 3D lung alveolar micro-architecture that SCLC grows in, we developed a microbead based tumor organoid that was scaled to a 96-well plate. We coated the beads first with ALFs from patients with no history of prior lung disease or lung cancer, as the main cell population (80% of total cells) using the ALF media and then introduced the H526 SCLC cells (20% of total cells) in the system using the co-culture media (mixture of ALF and SCLC media). Figure 1ai-iii demonstrates the growth of ALFs only on the microbead scaffold. Figure 1aiv-vi shows how the SCLC cells proliferated and invaded the ALF coated microbeads with a visible difference in phenotype compared to the 3D cultures that consisted only of ALFs. We used a vimentin promoter RFP reporter to visualize the ALFs and an EpCAM promoter GFP reporter to visualize the SCLC cells. Video S1 shows a 3D rendering of a whole SCLC-ALF co-culture organoid with the location of these cell types. To monitor the SCLC phenotype over time, we stained the organoids with the live cell permeant dye Calcein AM and imaged them at days 3-21 of culture (Figure 1b). Initially, we noticed SCLC cells surrounded the fibroblasts on the scaffold (Figure 1b, day 3) but between days 7 to 21 of culture, large SCLC cell clusters formed which took over the cultures and completely displaced the scaffolds showing an invasive tumor phenotype (Figure 1b, day 21). Video S2 shows the movement of SCLC cells surrounding the microbeads in the organoid microenvironment. We next calculated the tumor area in the 3D SCLC co-culture organoid using ImageJ, a Java based imaging program from the National Institutes of Health. We found a 9-fold increase in tumor area over days 3 to 21 in the SCLC organoid after which growth plateaued in the cultures (Figure 1c.), which may occur due to physical microenvironmental constraints.

**Figure 1.** Visual representation of the co-culture SCLC organoid model: a. Difference in healthy (ALF only) and cancer (SCLC-ALF) organoid phenotype after 72h. b. Change in SCLC-ALF organoid phenotype between culture day 3 and day 21 due to SSCLC cell growth. C. Tumor growth kinetics as observed by Calcein-AM (live cell dye)-stained organoid area measurement. D. Plot of Young’s modulus of whole organoid model shows the increase in organoid stiffness over time in culture.

Tissue mechanics, including tissue stiffness, provide physical cues that are a vital microenvironmental factor that can affect cell behavior. We measured the physical stiffness of the cells in our SCLC co-culture organoid over time in culture. We used JPK Nanowizard 4a Atomic Force Microscopy to show that the change in stiffness in terms of Young’s Modulus was increased about 5-fold from day 7 to day 21 of culture, as shown in Figure 1d. This progressive increase in cell stiffness may result from increasing cell-cell and cell-extracellular matrix (ECM) interactions as the tumor grows over time. Taken together, this SCLC-ALF co-cultured organoid recapitulates rapid 3D tumor growth, the invasive behavior of SCLC, and mechanobiological changes seen in SCLC. We therefore next sought to validate the SCLC organoid model for the production of neuroendocrine markers seen in SCLC tumors from patients.

### Neuroendocrine markers are expressed similarly in the SCLC organoid and in the SCLC patient tumor

SCLC is the most common form of neuroendocrine lung cancer and produces the classic neuroendocrine markers, Chromogranin A, synaptophysin, NCAM (CD56) and Calcitonin gene-related peptide (CGRP), which are commonly used as SCLC tumor markers [4, 9]. To further validate the co-cultured organoid’s pathophysiological characteristics, we tested the expression of CGRP, Chromogranin A, NCAM and Synaptophysin in the organoid at 14 days of culture and compared this expression with patient SCLC tumor sections, PDX tumors and classic 2D nonadherent SCLC cell line cultures. The PDX tumors and 2D cell cultures were developed from the same SCLC cell line as was used to generate the SCLC-ALF co-cultured organoids.

Immunostaining of a representative SCLC patient tumor demonstrated expression of CGRP, Chromogranin A, NCAM and Synaptophysin (Figure 2a). The PDX tumors (Figure 2b) and the SCLC cell line cultures (Figure 2c) both showed low expression of CGRP, Chromogranin A, NCAM and Synaptophysin whereas the SCLC-ALF co-cultured organoid model demonstrated similar expression of CGRP, Chromogranin A, NCAM and Synaptophysin as that of the SCLC patient tumor (Figure 2d). As the SCLC organoid model closely recapitulated the neuroendocrine expression of SCLC patient tumors, we next sought to examine the response of the SCLC organoid model to chemotherapy.

**Figure 2.** SCLC classic neuroendocrine biomarkers are expressed similarly in the SCLC co-culture organoid and in the SCLC patient tumor: CGRP, Chromogranin A, NCAM and Synaptophysin expression in a) patient tumor; b) patient-derived xenograft (PDX), c) SCLC cell line monoculture and d) SCLC co-culture organoid tumor. PDX, SCLC monoculture and SCLC co-culture organoid were developed from the same SCLC cell line (H526). The scale bar is 20 μm.

### Combining SCLC cells with lung fibroblasts more closely recapitulates SCLC tumor relapse after chemotherapy

As the SCLC-ALF co-cultured organoids demonstrate several key features of the disease, we tested their ability to model the relapse of SCLC after chemotherapy, which is the most challenging aspect of this tumor. For this we used the standard combination chemotherapy of Cisplatin and Etoposide at the half maximal inhibitory concentration (IC50), as described in the methods section. All IC50 values can be found in Supplementary Table 1. In order to better understand the role of ALFs in the SCLC cell’s response to chemotherapy, we also developed monoculture organoids containing only SCLC cells, cultured in monoculture media. Both mono- and co-cultured SCLC organoids were treated with the same doses of Cisplatin and Etoposide and monitored for cell survival. A brief workflow of this chemotherapy exposure experiment is shown in Figure 3a.

**Figure 3.** Combining SCLC cells with ALFs more closely recapitulates SCLC tumor relapse after chemotherapy: (a) experimental timeline. Visual representation of cell survival after chemotherapy as observed by (b) monoculture of SCLC cells and c) co-culture of SCLC cells and ALFs (scale bar: 500 μm). SCLC cell survival in the model plotted by (d) live cell fluorescence intensity measurement and (e) Cell-titre glo cell viability assay. ALFs in the model significantly increased secretion of the cytokines (f-g), chemokines (h-j) and MMPs (k-l) before (0 DPT) and after regrowth of cells (31 DPT) after chemotherapy.

We found that both the SCLC mono- and co-cultured organoid tumors had formed similarly by day 7 of culture. We then added Cisplatin and Etoposide (0 days post treatment (0 DPT)) at their combined IC50 dosage into the respective mono- and co-culture organoid media. After 72h, we removed the Cisplatin and Etoposide and monitored the organoids weekly for any change in live cell dye Calcein AM uptake (live cell fluorescence intensity, reflecting a change in live cell number) (Figure 3b). The Calcein stained image series shows the cell survival timeline observed in both the mono- and co-cultured SCLC organoids. In the monoculture SCLC organoid (Figure 3b), the SCLC tumor-like cell clusters are seen before chemotherapy (0 DPT) but by 10 DPT, there are almost no visible live cells until about 38 DPT when the SCLC cell clusters regrow. This is quantified by fluorescence intensity (Figure 3d). On the other hand, the co-culture SCLC organoid (Figure 3c) also showed a large reduction in visible live cells by 10 DPT but there was never a complete disappearance of live cells and large SCLC cell clusters were seen by day 31 post therapy. We further examined the mono- and co-culture SCLC organoids for cell viability using the Cell titre glo assay (Figure 3e). This quantification showed the same cell survival behavior as the fluorescence quantification from the Calcein intravital imaging. We also observed that the tumors co-cultured with ALFs showed significantly (P<0.0001) higher cell viability post chemotherapy, which could result from ALF resistance to chemotherapy and/or increased SCLC cell resistance in the presence of ALFs.

To further investigate this striking difference in cell survival post-chemotherapy between mono- and co-culture SCLC organoids, we examined the acute inflammatory factors secreted into the media that are typically associated with lung cancer progression, metastasis and angiogenesis [15–17]. Cell culture supernatants secreted by both types of SCLC organoids were collected at 0 DPT and 31 DPT and analyzed for secreted cytokines (interleukin (IL), IL-6, IL-8), chemokines (growth regulated alpha protein (GRO-α), monocyte chemoattractant protein-1 (MCP-1)), vascular endothelial growth factor (VEGF-A)), and matrix metalloproteinases (MMP1, MMP2).

For both the 0 DPT and 31 DPT timepoints, all the inflammatory secreted factors tested were present in significantly (P<0.0001) higher amounts in the co-culture SCLC model than in the monoculture SCLC model (Figure 3f-l). One explanation for this is that most of the secreted factors come from the ALFs, and that this allows for the more robust survival of cells after chemotherapy in the SCLC co-culture organoids. But as the SCLC monoculture organoids are capable of producing all of these secreted factors at low level, we next asked whether the ALFs could induce the SCLC cells to produce more of these factors, which could promote SCLC progression in the SCLC co-culture organoids. We, therefore, investigated whether direct cell-cell interactions between the ALF and SCLC cells are needed for cell survival after chemotherapy in the co-culture SCLC organoids or whether this is due to a paracrine effect.

### Secreted paracrine factors from lung fibroblasts drive the regrowth of SCLC cells in our tumor organoid model

In order to better understand whether regrowth of SCLC cells after chemotherapy may be driven by paracrine factors from healthy adjacent fibroblasts, we used monoculture organoids cultured in either ALF-conditioned media or in usual monoculture media, and treated with Cisplatin and Etoposide, as described previously. In both of the cases, the monoculture organoids were cultured for 38 days after chemotherapy and the inflammatory factors secreted in both conditions were examined from time points before treatment (0 DPT) and at 31 days after treatment (31 DPT). The experimental details and timelines are shown in Figure 4a. We used Calcein AM intravital dye imaging to demonstrate the monoculture organoid before drug treatment (0 DPT) and regrowth of SCLC cells in the organoid mimicking tumor relapse at 31 DPT. It is evident from Figure 4b that the live cell regrowth in monoculture SCLC organoids with ALF-conditioned media was similar to the co-culture SCLC organoids in co-culture media in Figure 3c and more robust in both of these conditions than monoculture SCLC organoids grown in monoculture media only (Figure 3b).

**Figure 4.** Effect of ALF-conditioned media on SCLC relapse: (a) experimental timeline. (b) visual representation of a SCLC monoculture organoid grown in ALF-conditioned media before chemotherapy and after cell regrowth (scale bar: 500 μm). ALF-conditioned media significantly increased secretion of the chemokines, cytokines and MMPs before and after chemotherapy (c-i). (j) ALFs and SCLC cells are not closely opposed to each other when the SCLC cells regrow after chemotherapy (scale bar: 200 μm).

As elucidated in Figure 4c-i, monoculture SCLC organoids in ALF-conditioned media contained higher levels of the secreted inflammatory factors than monoculture SCLC organoids in monoculture media only. Because the same batch of ALF-conditioned media was used throughout the whole experiment, we next compared levels of each inflammatory factors secreted by the monoculture organoids in ALF-conditioned media at 0 and 31 DPT. Comparison of each factor before and after SCLC cell regrowth revealed that GRO-α, IL-8 and MMP-1 had significantly (P<0.001) higher expression in the media with SCLC cell recurrence than in the original tumor before chemotherapy. This suggests that the ALF-conditioned media induced the SCLC cells to secrete more GRO-α, IL-8, and MMP-1 and these factors all play key roles in angiogenesis and tumor metastasis [19–22]. Interestingly, the monoculture organoids cultured in monoculture media showed a trend in secreting more GRO-α, IL-8, MMP-1 and MMP-2 at DPT 31 than at DPT 0, but to a much smaller extent than the co-cultures, showing that SCLC cells secrete more of these factors when they survive after chemotherapy even in the absence of ALFs.

We further examined the location of the ALF and SCLC cells in our co-culture model during chemotherapy and regrowth (0-38 DPT) to examine the proximity of these cells for cell-cell interactions. Vimentin-expressing ALFs (red) and EpCAM-expressing SCLC cells (green) were initially patterned in close opposition to each other (0 DPT) surrounding the bead scaffolds with SCLC cells being the dominant population (Figure 4j). Post-chemotherapy (10 DPT), disruption of the SCLC organoid structures takes place keeping ALFs alive with noticeably, a few surviving SCLC cells. With time in culture, (24-38 DPT), the SCLC population takes over the culture again and the fibroblasts are found on the periphery of the tumor regions. Therefore, at least in the context of cell survival after chemotherapy, the paracrine effects from ALFs seem to contribute more to SCLC cell survival after chemotherapy than direct SCLC cell-ALF interactions.

## Discussion

Here, we have developed a scaffold-based 3D co-culture SCLC organoid model using a top-down approach. The addition of SCLC cells to the ALFs in the lung organoid microarchitecture reveals the phenotypic transformation and kinetics of *in vitro* tumor formation. The immunofluorescent staining of the SCLC co-culture organoids reveals the expression of neuroendocrine biomarkers that are more highly expressed than the PDX models from the same cell line. The SCLC organoid co-culture model was also used to assess microenvironmental parameters like mechanobiological changes over time in culture. Stiffening of tumors is reported to be a sign of tumor microenvironment remodeling with tumor cell growth, displacement of host tissue, and cancer cell invasion of surrounding tissues [14, 23–25]. We observed increasing tumor stiffness and displacement of the scaffold with time in culture suggesting that this co-culture model could be useful for analyzing the mechanobiological changes that occur with cancer progression.

Relevant SCLC models that mimic the tumor relapse seen in patients are challenging to develop because the tumor microenvironment is key for developing this phenotype. We generated a scalable 3D SCLC co-culture model in the dish that can be used to study this biology and can be used for high throughput drug screening (HTS). We used the SCLC organoid model to study the role of ALFs in SCLC co-culture organoids and the model lends itself to the addition of other cell types to further understand the role of the microenvironment in disease progression and relapse. Our model was also able to distinguish between the effects of direct cell-cell interactions and paracrine signaling between ALFs and SCLC cells on tumor cell survival and this has implications for therapeutic development.

Taken together, our study shows that we have built a powerful new tool for studying SCLC that is modular and allows the incorporation of multiple cell types and microenvironmental factors that influence tumor behavior. Currently, there is no *in vitro* model of human SCLC that phenocopies the tumor microenvironment and demonstrates its effects on chemoresistance. Our SCLC organoid model can be used to answer in-depth biological questions on SCLC tumor development, progression, and relapse and is scalable for HTS.

## Materials and Methods

### Scaffold generation and functionalization

We prepared alginate microbead scaffolds (average diameter 100 um) using a hydrostatic droplet generator. We used 3% sodium alginate solution as the biopolymer base and 100mM Barium chloride solution as the crosslinking agent to prepare the beads. Once the beads were made, we functionalized them with a two-step coating of collagen I (Corning 354249) and dopamine (Sigma H8502) as extracellular matrix (ECM) to prepare them for cell culture. The detailed method for alginate bead generation and functionalization is described in Wilkinson et al [22]

### SCLC and ALF cell culture and media preparation

Human SCLC cell lines H526 (NCI-H526) (CRL-5811) were purchased from ATCC. After receiving the cells, they were cultured in standard tissue culture flasks (Genesee scientific) in the SCLC media [RPMI 1640 (Gibco),10% fetal bovine serum (ATCC), 0.2% primocin] and cell passages <10 were used for this study.

Human primary adult lung fibroblasts (ALF) were isolated from distal lung tissue from a de-identified healthy donor (65-year-old, male, Caucasian, non-smoker, non-alcoholic) procured from the International Institute for the Advancement of Medicine (IIAM). Human lung tissue was procured under the UCLA approved IRB protocol #16-000742 which approved all experimental protocols. Methods were carried out in accordance with all relevant guidelines and regulations. The distal tissue was cut into 1cm×1cm pieces and kept in 6-well plates, submerged in ALF media (DMEM+F12, 10% FBS, 1% non-essential amino acids, 1% glutamax) for 3-4 weeks to allow the fibroblasts to “crawl out” of the tissue and form an adherent mono layer in the well. The “crawled-out” populations were dissociated with TrypLE (Thermofisher 12605036), cultured in cell culture flasks and passages <5 were used for this study.

SCLC media is referred to as monoculture media and the combination of SCLC and ALF media (4:1) is referred to as co-culture media throughout this manuscript.

For the ALF-conditioned media preparation, a subculture of ALFs was cultured in ALF media without serum for ~48h in a separate flask until ~80% confluency was achieved. The culture supernatant was then collected, filtered and used for the entirety of the experiment.

### Bioreactor setting, 3D model formation and loading into 96-well plate

To develop the 3D model, we used a high aspect ratio vessel (HARV) bioreactor vessel (model: RCCS-4H; Synthecon, Houston, Texas) of 2 ml volume and added 0.5 ml of functionalized microbeads and 1.5 ml of media containing a total of 1 million cells. The vessel was screwed into the bioreactor base and the beads and the cells allowed to settle. After sedimentation, the bioreactor was powered on to 4 rpm.

For the co-culture model, ALFs and SCLC cells were mixed in a 4:1 ratio and the co-culture media was used. To mimic the natural cellular layering, the ALFs and beads were added first in the bioreactor. Once the beads were coated with ALFs (~6h), the SCLC cells were added in the bioreactor and rotated until a uniform 3D structure was formed (~48h).

For the monoculture model (SCLC cells only), 1 million SCLC cells with 0.5 ml beads and 1.5 ml monoculture media were rotated for 48h.

For the ALF-only model (Figure 1a), 1 million ALFs with 0.5 ml beads and 1.5 ml ALF media were rotated for 48h.

For all cellular combinations, after 48h, the cell-coated bead solution was aliquoted 100μl per well in a glass-bottom 96-well plate (Cellvis P96-1.5H-N) with the help of a multichannel pipettor. The 96-well plate was then briefly centrifuged (1000g, 2 min) to settle the cells/ beads at the bottom of the plate and an additional 150μl media was added to each well. The plate was then kept inside an incubator (37°C, 5% CO_2_, 95%RH) and monitored for the formation of self-organized 3D structures. Within the next 72h, the fully-formed 3D models with micro-alveolar structures were observed in each well.

From one bioreactor of 2ml capacity, 20 such models were formed. The number and capacity of the bioreactors were varied as required. For a full 96-well plate, we used 5×2ml bioreactors or 1×10ml bioreactor.

### Atomic force microscopic analysis

For tumor stiffness measurements, 7-, 14- and 21-day old live 3D co-culture models were transferred to 35mm fluorodishes (WPI, FD35-100) containing phosphate-buffered saline (PBS) buffer and atomic force microscopy was performed at 37°C in PeakForce Tapping mode using JPK Nanowizerd 4A (Bruker Nano Surface, CA, USA). We used a PeakForce Quantitative Nanomechanics-Live Cell (PFQNM-LC) probe (Bruker AFM probes, CA, USA) with a silicon tip [length=54μm; radius=4.5μm; frequency=45kHz; Spring constant=0.1N/m], specially optimized for soft biological samples. During measurements, multiple (n ≥3) tumor locations were selected and an area of 100μm×100μm was scanned in each location. Force-distance curves were recorded to obtain tumor stiffness. Data analysis was done in JPKSPM Data processing software (version 6)(Bruker, USA). For calculating Young’s modulus, force-distance curves were converted to force-separation curves and Hertz-Sneddon model was chosen during model fitting.

### Chemotherapy treatment and relapse study

For the chemotherapy treatment, Cisplatin (Tocris, Batch No: 5B/266434) and Etoposide (Sigma, Lot # 099M4892V) were used. The drugs were added to the wells containing 7-day old SCLC tumors at a concentration equal to their respective half-maximal inhibitory concentration (IC50). To calculate the IC50, the tumors were treated with cisplatin and etoposide, singly and in combination, in a range of 0-100μM, for 72h. Upon reaching the end point, cell viability was determined via Cell-Titre-Glo assay (Promega) according to manufacturer’s protocol for viability determination. From the concentration and corresponding viability values, IC50 was calculated using Graphpad Prism (version 9) software. All treatments were done in triplicate.

To study cancer relapse after chemotherapy treatment, we measured cell viability by image analysis and using Cell-Titre-Glo kit (Promega) to detect metabolically active cells. All readings were taken in triplicate at 7-day intervals for a duration of 0 to 38 post-drug treatment days.

### Imaging of live organoids and image analysis

For video 1, the ALFs were tagged with a vimentin promoter RFP reporter and SCLCs were tagged with an EpCAM promoter GFP reporter and the 3D rendering video was captured with a Leica Thunder confocal microscope.

For other live cell imaging, the live organoids were stained with Calcein-AM (Thermofisher, C3099) a GFP fluorescent live cell labeling dye at 1:1000 dilution, and imaged in a Zeiss Inverted Phase Contrast Fluorescence Microscope (Axiovert 40 CFL) at regular intervals (as per experimental design). For each time point and each treatment, experiments were done in triplicate. For image analysis, images of the same set of wells were captured weekly to monitor for change over time. Every time, fresh Calcein AM dye was added and any residual dye was washed off with media after the imaging was completed.

We used ImageJ (National Institute of Health, USA) software to analyze tumor area. For a particular image, the GFP zones were selected and the inbuild area measurement tool of the software was used to calculate area [26]. The scale was converted from pixel to equivalent micron using the actual scale of the image.

For the intensity analysis of an image, we first computed a threshold to separate out the background from the foreground component of the image. In particular, we used the popular Otsu’s method [27] to compute the intensity threshold *I_th_*. Otsu’s method instinctively performs clustering-based thresholding that assumes two classes of pixels backing bi-modal histogram (foreground and background pixels). For each image, we then compute the binary mask of dimension the same as that of the image, with logical ones corresponding to the pixel locations having value > *I_th_*. To compute the positive intensity, we then evaluate the mean of the masked image. Similarly, for negative intensity evaluation we compute the intensity of the complementary masked image. To expedite the intensity computation and allow automated intensity evaluation for an entire image batch, we developed a script in Python language. Additionally, the presented automated intensity analysis process may significantly reduce human error associated with image-wise manual intensity evaluation.

We used Python language (version 3.7) to evaluate the intensity for the collected image set. The images were saved in grey scale .tif format with a resolution of 1269 x 972.

The example of intensity analysis is shown in Supplementary Figure S1b.

### Immunofluorescence staining and imaging of fixed organoid

For whole-mount staining, the organoids were first fixed using 4% paraformaldehyde (Thermo Fisher) for 30 minutes at room temperature and then permeabilized using 0.1% TritonX-100 (Sigma-Aldrich) in in Tris-buffered saline (TBS) for 15 minutes. After blocking in DAKO (Agilent) for 1h, organoids were incubated with primary antibodies overnight at 4°C. Next day, after triple washing, organoids were incubated in secondary antibodies (ThermoFisher) along with 4’,6-diamidino-2-phenylindole (DAPI) for 1h at room temp. The following primary antibodies were used: mouse anti-vimentin (Abcam), rabbit anti-calcitonin gene related peptide (CGRP) (Millipore Sigma), rabbit anti-EpCAM (Abcam), rabbit anti-synaptophysin (Abcam), mouse anti-chromogranin A (Proteintech), NCAM1 (Cell Signaling Technology). Confocal imaging was performed using Zeiss LSM 880.

### Statistical analysis

All data were compiled from three or more independent replicates for each experimental condition. Data comparisons were performed using unpaired Student’s *t*-tests, multiple comparisons tests and correlation analysis using 2-way ANOVA using the GraphPad Prism software (version 9).

## Acknowledgements

This work was supported by the American Fund for Alternatives to Animal Research (AFAAR) (CS), the Keck Foundation (BNG), the Tobacco-Related Disease Research Program grant T31DT1684 (DWS), the NIH/NCI Grant R01CA208303 (BNG), the Tobacco Related Disease Research Program (TRDRP) High Impact Pilot Research Award (HIPRA) 26IP-0036 (BNG), the TRDRP HIRA 29IP-0597 (BNG), The UCLA Jonsson Comprehensive Cancer Center, (JCCC) STOP Cancer Award (BNG), the Ablon Research Scholars Award (BNG). Statistical analyses were supported by the NIH/National Center for Advancing Translational Science (NCATS) UCLA CTSI Grant Number UL1TR000124. We thank the staff of the UCLA BSCRC Microscopy Core, UCLA CNSI NPC core and the UCLA Translational Pathology Core Laboratory. Figures 3a and 4a were created with BioRender.com.

## Authors’ Contributions

C.S. and B.N.G. conceived of the study. C.S. designed experiments. C.S., C.K., S.K., D.W., J.Y., D.W.S., and L.K.M., performed experiments and acquired data. C.S. and B.N.G. interpreted data. C.S. and B.N.G. wrote the manuscript. C.S., C.K., S.K., D.W., D.W.S., J.Y., L.K.M., T.R. and B.N.G. provided administrative, technical, or material support.

## Data availability statement

Datasets generated in the study are available from the corresponding author on reasonable request.

## Figure legends

**Video 1**: Biorendering of a co-culture SCLC organoid with small cell lung cancer (SCLC) cells (GFP tagged for Epcam reporter) and adult lung fibroblast (RFP tagged for Vimentin reporter).

**Video 2**: Movement of cancer cells surrounding the microbeads in the co-culture SCLC organoid model.

