## Supplementary Figures and Table for "Small cell lung cancer co-culture organoids provide insights into cancer cell survival after chemotherapy"

**Supplementary Table 1**

IC50 values of Cisplatin and Etoposide singly and in combination

| | Cisplatin (µM) | Etoposide (µM) | Combined (µM) |
|---|---|---|---|
| H526 | 7.97±1.26 | 2.1±0.83 | 3.68±0.44 |

**Supplementary Figure legends:**

S1. Monocultures of SCLC cell lines in the dish showed significantly less viable cells after chemotherapy treatment than the co-culture SCLC organoid.

S2. Example of thresholding accuracy during automated image intensity measurement

**Supplementary Figure 1**

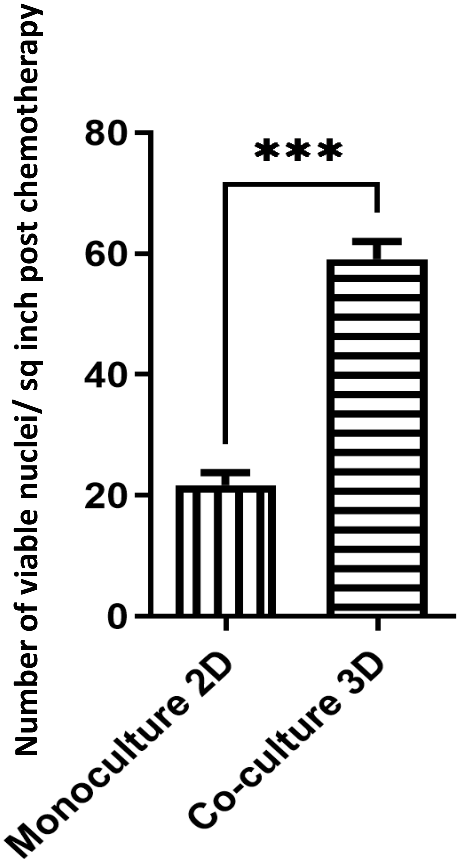

**Supplementary Figure 2**

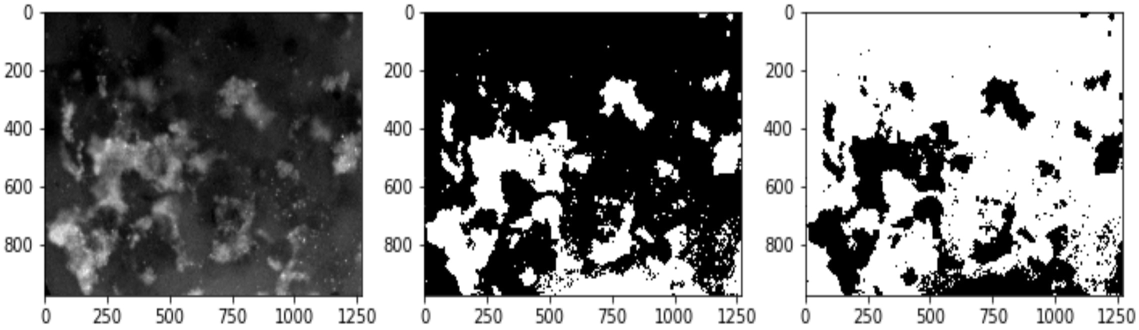
